# Isolation and genomic characterization of the tick-borne relapsing fever spirochete, *Borrelia turicatae*, from ticks collected in a peridomestic setting of Camayeca, Mexico

**DOI:** 10.1101/2023.08.01.551332

**Authors:** Edwin Vázquez-Guerrero, Alexander R. Kneubehl, Patricio Pellegrini-Hernández, José Luis González-Quiroz, María Lilia Domínguez-López, Aparna Krishnavajhala, Paulina Estrada-de los Santos, J. Antonio Ibarra, Job E. Lopez

## Abstract

Surveillance studies were implemented in Sinaloa, Mexico to determine the circulation of tick-borne relapsing fever spirochetes. Argasid ticks were collected from a human dwelling in the village of Camayeca and spirochetes were isolated. Genomic analysis indicated that *Borrelia turicatae* is a threat to those living in resource limited settings.

**Article Summary Line:** We report the collection of argasid ticks from a peridomestic setting in Mexico and the isolation of *Borrelia turicatae*; increased surveillance efforts are needed on this overlooked vector-borne pathogen.

## Introduction

Tick-borne relapsing fever (TBRF) spirochetes are neglected pathogens in Mexico and likely misdiagnosed. For example, *Borrelia turicatae* presents with nonspecific symptoms like irregular fevers, meningitis, rigors, nausea, and vomiting (1). The neurological manifestation of disease can be misleading resulting in a misdiagnosis of Lyme disease. Furthermore, the use of nonspecific serological tests further complicates an accurate diagnosis of TBRF. Whole spirochete lysates of *Borreliella* (*Borrelia*) *burgdorferi* have been used for ELISA and immunoblotting (2, 3), yet it is known that serological cross reactivity occurs regardless of whether patients are infected with Lyme-causing or TBRF spirochetes (4).

Argasid ticks transmit most species of TBRF spirochetes, and their life cycle further complicates a clear understanding of the disease’s epidemiology. Argasids in the genus *Ornithodoros* are cavity dwelling rapid feeders that are rarely found attached to the host. In a case study, people reported being bitten by insects during their sleep (5). An investigation of the home identified *Ornithodoros puertoricensis* under floor tiles and within cracks of windowsills (5). These findings indicated that once introduced into the dwelling, the occupants were the primary blood source for the ticks.

With the need to better understand the distribution of TBRF spirochetes and their vectors in Latin America, we initiated efforts to collect argasid ticks from peridomestic settings of Mexico. We describe the identification of *Ornithodoros turicata* from the village of Camayeca in Sinaloa, Mexico. The ticks were determined to be infected by feeding them on a laboratory mouse. TBRF spirochetes were isolated from mouse blood and genomic analysis speciated the spirochetes. Our results identified an endemic focus of *B. turicatae* in the village of Camayeca, Mexico.

### The study

In March 2022 we collected argasid ticks in peridomestic settings of Sinaloa, Mexico. In the village of Camayeca (Figure 1A), we sampled five burrows using an aspirator or dry ice as a source of carbon dioxide to lure ticks. In the human dwelling where ticks were collected (Figure 1B), we aspirated the dirt at the base of the home (Figure 1C). We collected three adults and 19 nymphs. At this location we noted ground squirrel activity around the burrows.

**Figure 1.**
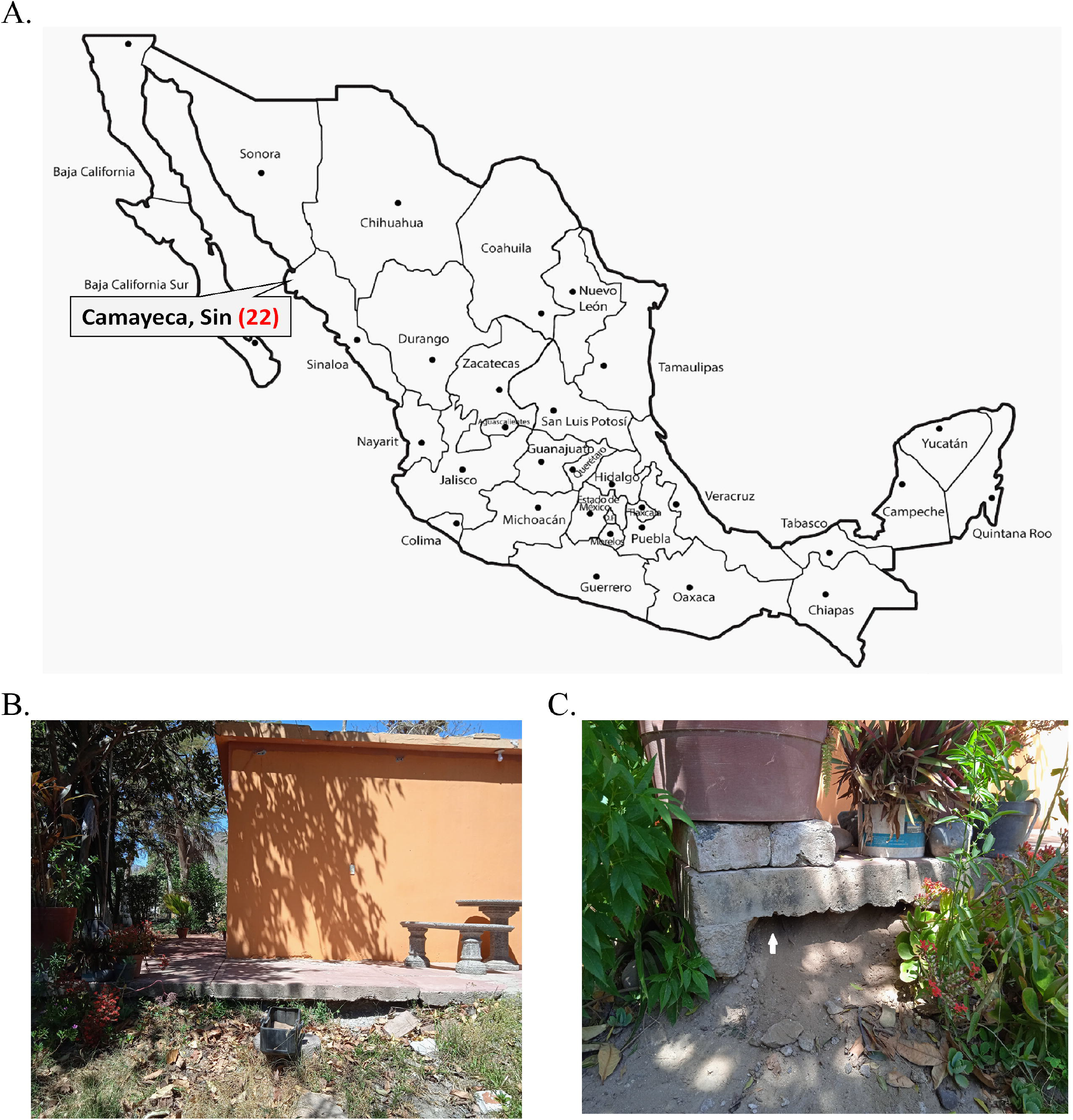
Collection of *Ornithodoros turicata* in village of Camayeca, Mexico. Shown is a map of Mexico and the state of Sinaloa where 22 ticks were collected (A). Collection efforts were focused in peridomestic settings (B), and ticks were aspirated from the base of a human dwelling (C). The white arrow in (C) points to where ticks were collected.

In the laboratory, we speciated ticks using microscopy and through sequencing a portion of the 16S mitochondrial gene. Morphological characterization of nymphs and adults indicated that they were *Ornithodoros turicata*. Total DNA was also extracted from three nymphs using the Qiagen DNeasy Blood and Tissue kit (Qiagen, Germantown, MD, USA) following the manufacturer’s protocol. We amplified ∼475 nucleotides of the 16S mitochondrial rRNA gene using Tm16S+1 and Tm16S-1 primers (6). The amplicons were Sanger sequenced and the data trimmed using ChromasPro v. 2.1.5 (Technelysium Pty Ltd). We performed a BLASTN analysis on the NCBI website, which indicated 99.1% nucleotide identity to *O. turicata*. Sequences were deposited to GenBank under accession numbers: OR189376 - OR189378.

We determined whether the remaining ticks were infected with TBRF spirochetes by allowing them to feed on a BALB/c mouse in accordance with the Institutional Animal Care and Use Committee (protocol # ZOO-001-2022). Daily blood samples were collected from the mouse and Giemsa staining was performed to visualize spirochetes. Seven days after feeding ticks the mouse was exsanguinated and whole blood was centrifuged at 500 g for 5 min. Plasma was removed and centrifuged again at 5,000 g for 10 min. The pellet was resuspended in 1 ml of Barbour-Stoenner-Kelly (BSK)-R media and cultured in a total of 4 ml at 35 °C (7). Eight days later an aliquot of the culture was placed on a glass slide, air dried, and Giemsa stained. We visualized numerous spirochetes on the slide (Figure 2A). The isolate was designated CAM-1 and glycerol stocks were generated.

**Figure 2.**
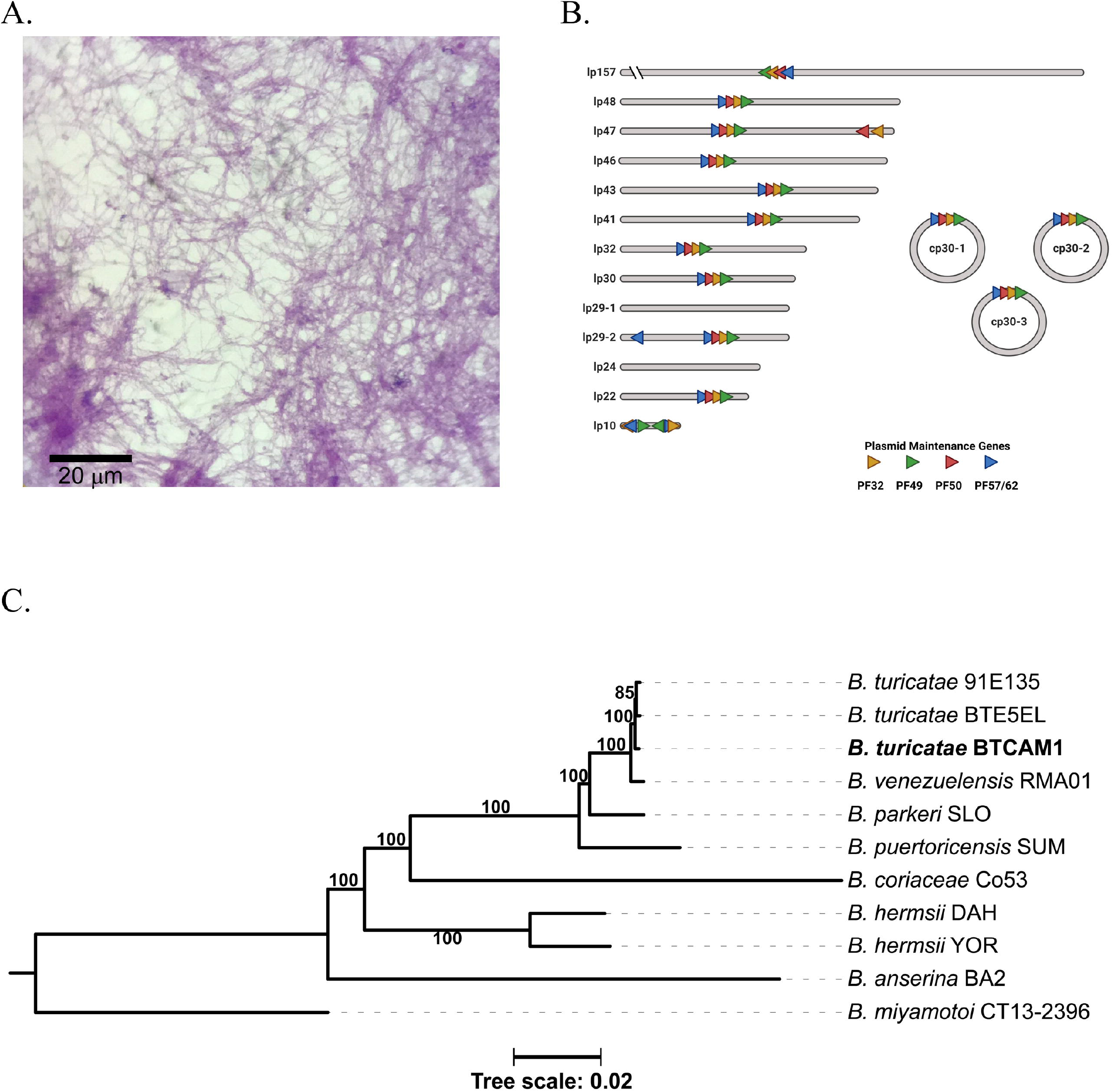
Isolation and genetic characterization of TBRR *Borrelia*. Spirochetes were isolated from murine blood in culture medium (A). Genome sequencing and assembly generated the plasmid repertoire of the bacteria (B). Plasmids were designated as linear (lp) or circular (cp) and by their respective size to the nearest kilobase. The plasmid family (PF) partitioning genes are shown in each plasmid as orange, green, red, and blue triangles. A maximum likelihood species tree was performed in a phylogenomic analysis of CAM1 and grouped the spirochete with *B. turicatae* (C). The tree was generated with an edge-linked proportional partition model with 1,000 ultra-fast bootstraps.

We sequenced the CAM-1 isolate to determine the species and number of plasmids that it harbored. Genomic DNA was isolated and pulsed field electrophoreses was performed to determine DNA quality, as previously described (8). Long-read sequencing was performed with the Oxford Nanopore Technologies (ONT) Mk1B platform with the SQK-RBK110.96 library preparation kit and R9.4.1 flow cell. Short-read sequences were generated by Microbial Genome Sequencing Center (MiGS Center, USA) using an Illumina 2×150 library preparation kit. We produced a plasmid-resolved genome assembly by using short-reads to polish the long-read data, as previously reported (9). The mean ONT coverage was 439x and the mean Illumina coverage was 236x. Using a previously established approach (9), completeness and QV scores (based on the Phred scale) were 99.89% and Q53.82, respectively. The assembly was annotated with NCBI’s Prokaryotic Genome Annotation Pipeline and submitted to NCBI’s GenBank (accession numbers CP129306-CP129322). The chromosome was 925,885 bp. There were 16 plasmids ranging from 10,351 to 156,755 bp, and three were circular (Figure 2B). The phylogenomic analysis used concatenated sequences from 650 core genes, which encompassed 720,532 nucleotides. This grouped the CAM-1 isolate with *B. turicatae* (Figure 2C).

## Conclusion

Until our work, an isolate of *B. turicatae* has been absent from Latin America. In this current study, we collected ticks from a human dwelling in the village of Camayeca. We identified the ticks as *O. turicata* and determined that they were infected by feeding them on a mouse. Upon isolating spirochetes from the animal’s blood, a phylogenomic analysis indicated that the bacterium was the pathogenic spirochete, *B. turicatae*.

While *O. turicata* was originally described in Mexico in 1876 by Alfredo Dugès (10), the tick was not implicated as a vector of TBRF spirochetes until the 1930s. In 1933, Brumpt and colleagues detected spirochetes in *O. turicata* collected from Austin, Texas, and he later confirmed that the tick species could transmit *B. turicatae* (11). At the same time, Pilz and Mooser were detecting human cases of relapsing fever in the city of Aguascalientes, Mexico (12). Their work showed that *O. turicata* was in the region and implicated the tick species as the vector. Our identification of *B. turicatae* in Camayeca, which is over 1,000 km from the city of Aguascalientes, broadens the range of TBRF spirochetes in the country.

Our findings indicate that updates are needed for species distribution models of *O. turicata*. For example, a maximum entropy species distribution model analysis predicted suitable habitat of *O. turicata* and used a stringent definition to generate georeferenced data points (13). While new regions of northern Mexico were predicted to have habitat for *O. turicata*, there was low probability of suitable habitat in other regions of the country. In addition to collecting ticks from Camayeca, Mexico, we have also recovered *O. turicata* from the city of Aguascalientes, Mexico (manuscript in review). The city of Aguascalientes is in the middle of the country and is considered a temperate environment ∼1,900 meters in elevation while Camayeca is an arid desert region at ∼150 meters above sea level. The environmental differences between these two cities indicates wider habitat suitability for *O. turicata* than what was previously predicted.

Identifying infected *O. turicata* in a peridomestic setting suggests that the disease is likely under reported in Mexico. In support of this, retrospective serodiagnostic studies indicate human exposure to TBRF spirochetes in populations originally diagnosed with fever of unknown origin (14). That work and our current findings indicate the importance of understanding the distribution and ecology of *O. turicata* and other argasid ticks of human importance in Mexico.

## Acknowledgments

We thank Dr. Miguel Medina-Cota for putting us in contact with our collaborator in Sinaloa.

## Funding

This work was supported by funds provided to JEL from the National School of Tropical Medicine at Baylor College of Medicine and by funds to JAI from Secretaría de Investigación y Posgrado-IPN (20230850).

## About the Author

Dr. Vázquez-Guerrero is an infectious disease specialist in Mexico. His areas of interest are acarology and infectious diseases.

## Notes

### Competing Interest Statement

The authors have declared no competing interest.

## References

1. Cadavid D, Barbour AG. Neuroborreliosis during relapsing fever: review of the clinical manifestations, pathology, and treatment of infections in humans and experimental animals. Clinical Infect Dis. 1998 Jan;26(1):151–64.

2. Gordillo-Perez G, Solorzano F, Cervantes-Castillo A, Sanchez-Vaca G, Garcia-Ramirez R, Diaz AM, et al. Lyme neuroborreliosis is a severe and frequent neurological disease in Mexico. Arch Med Res. 2018 Aug;49(6):399–404.

3. Gordillo-Perez G, Garcia-Juarez I, Solorzano-Santos F, Corrales-Zuniga L, Munoz-Hernandez O, Torres-Lopez J. Serological evidence of Borrelia burgdorferi infection in Mexican patients with facial palsy. Rev Invest Clin. 2017 Nov-Dec;69(6):344–8.

4. Lopez JE, Schrumpf ME, Nagarajan V, Raffel SJ, McCoy BN, Schwan TG. A novel surface antigen of relapsing fever spirochetes can discriminate between relapsing fever and Lyme borreliosis. Clin Vaccine Immunol. 2010 Apr;17(4):564–71.

5. Bermudez SE, Castillo E, Pohlenz TD, Kneubehl A, Krishnavajhala A, Dominguez L, et al. New records of Ornithodoros puertoricensis Fox 1947 (Ixodida: Argasidae) parasitizing humans in rural and urban dwellings, Panama. Ticks Tick Borne Dis. 2017 Feb 05.

6. Black WCt, Piesman J. Phylogeny of hard- and soft-tick taxa (Acari: Ixodida) based on mitochondrial 16S rDNA sequences. Proc Natl Acad Sci U S A. 1994 Oct 11;91(21):10034–8.

7. Replogle AJ, Sexton C, Young J, Kingry LC, Schriefer ME, Dolan M, et al. Isolation of Borrelia miyamotoi and other Borreliae using a modified BSK medium. Sci Rep. 2021 Jan 21;11(1):1926.

8. Simpson WJ, Garon CF, Schwan TG. Analysis of supercoiled circular plasmids in infectious and non-infectious Borrelia burgdorferi. Microbial Pathogenesis. 1990;8:109–18.

9. Kneubehl AR, Krishnavajhala A, Leal SM, Replogle AJ, Kingry LC, Bermudez SE, et al. Comparative genomics of the Western Hemisphere soft tick-borne relapsing fever borreliae highlights extensive plasmid diversity. BMC Genomics. 2022 May 31;23(1):410.

10. Dugès A. Turicata de Guanajuato. Artículo en el periódico “El Repertorio” de Guanajuato. 1876;Sect. 11–2.

11. Brumpt E, Brumpt LC. Identite du spirochete des fievres recurrentes a tiques des plateaux mexicains et du Spirochaeta turicatae agent de la fievre recurrente sporadique des Etats-Unis. Ann Parasitol Hum Compar. 1939;17:287–98.

12. Pilz H, Mooser H. La fiebre recurrente en Aguascalientes. Bol Inst Hig México. 1936;2:295–300.

13. Donaldson TG, Perez de Leon AA, Li AI, Castro-Arellano I, Wozniak E, Boyle WK, et al. Assessment of the geographic distribution of Ornithodoros turicata (Argasidae): Climate variation and host diversity. PLoS Neg Trop Dis. 2016 Feb;10(2):e0004383.

14. Vazquez-Guerrero E, Gordillo-Perez G, Rios-Sarabia N, Lopez JE, Ibarra JA. Case Report: Exposure to relapsing fever group borreliae in patients with undifferentiated febrile illness in Mexico. Am J Trop Med Hyg. 2023 Mar 1;108(3):510–2.

